# Fabrication of silver doped nano hydroxyapatite-carrageenan hydrogels for articular cartilage applications

**DOI:** 10.1101/2020.12.31.424664

**Authors:** Subhasmita Swain, Tae Yub Kwon, Tapash R. Rautray

## Abstract

It can be found from the results that nano hydroxyapatite- silver -3.0 wt% carageenan (nHA-Ag-CG3.0) improved the mechanical properties of the as-formed hydrogel scaffold after incorporation of higher CG concentration. The Young’s modulus of hydroxyapatite- silver - 1.5wt% carageenan (nHA-Ag-CG1.5) was found to be 0.36 ± 0.07 MPa that increased in case of nHA-Ag-CG3.0 demonstrating better interfacial compatibility of their matrix with respect to the reinforcement. This increase in reinforcement concentration resulted in higher stiffness that dissipated energy. The higher swelling ratio is envisaged to induce better cell adhesion and proliferation. The biodegradability test was performed in phosphate buffered saline at body temperature for 3 weeks. The biodegradability rate of nHA-Ag-CG1.5 was found to be equivalent to nHA-Ag-CG3.0 hydrogels at day 7 while it increased faster in nHA-Ag-CG3.0 on days 14 and 21 that may be ascribed to the possible interaction of nHA and Ag with their CG matrix. The bacterial cell viability of *Staphylococcus aureus* (*S. aureus)* was performed after 10 h, 20 h and 30 h of culture. The nHA-Ag-CG1.5 exhibited restrained growth of *S. aureus* as compared to nHA-Ag-CG3.0 and these results were validated by CLSM analysis. Hence, nHA-Ag-CG3.0 may be considered to have more cytocompatibility than nHA-Ag-CG 1.5.

## 1. Introduction

The replacement of bone tissues caused due to accident or disease has become a new challenge in the field of biomaterials science. Therefore, fabricating a novel biomaterial having outstanding bone compatible properties that would replace the currently used biomaterials with many drawbacks has become vital in the field of tissue engineering. With the use of either physical or chemical crosslinking, a network of 3D hydrophilic structure termed as scaffolds, can be fabricated which have promising physical and biological properties including high water retention, favorable biocompatibility and lowest inflammation capability equivalent to natural functional tissues [1-4].

Production of scaffolds from biocompatible materials has recently been getting serious attention in biomedical engineering since they offer bio-physico-chemical pathways of increasing and restoring natural tissues. A 3D framework that mainly operates as a supporting structure for body cells to adhere and proliferate thus permitting them to grow and achieve their functions [5, 6]. As soon as the cells start to experience proliferation, degradation of the scaffold initiate and are slowly absorbed by the body. Gradually the scaffold gives space to the cells for more regeneration and the new tissues formed organize to take the total shape of scaffold. So as to facilitate regeneration of new tissues, some biomimetic and structural requirements are taken into account. A biomedically ideal scaffold must be biocompatible having interconnected pore channels so as to allow cell migration, supply oxygen and nutrients. The scaffold would promote cell adhesion, proliferation, differentiation and have controlled biodegradable rate having adequate mechanical strength to act as the support system for the cells [7-9].

HA is the primary inorganic constituent of bone and teeth that can be used as a potential reinforcement in hydrogels for bone tissue engineering applications. HA has long been widely used as bone substitute materials. In fact, because of its high osteoconductivity, excellent biocompatibility and bioactivity, HA can be extensively used as a guided bone regeneration material in spite of its low degradation rate and fragility [10-12].

Because of its slow degradation rate, application of HA as bioresorbable material is limited. To overcome this drawback, other calcium phosphate materials with higher solubility have been investigated to be used as bone-substitutes. Additional components such as Sr, Mg, Ag, Cu, Zn etc. are added to them for improving and inducing new properties [13, 14].

CG is a derivative of a marine algae called ‘Rhodophyceae’ and is a high molecular weight anionic heteropolysaccharide. It is a low cost, non-toxic, biocompatible and hydrophilic marine based linear polysaccharide. Moreover, the sulfate groups present in it has the potential to mimick glycosaminoglycans (GAGs) which are the negatively charged macromolecules [15, 16]. CG in combination with HA would be a potential candidate in orthopaedic applications.

Furthermore, the gel strength has been found to be higher in case of less sulfate concentration in carrageenan and the least ester sulfate is available in κ-carrageenan that can be most appropriate for applications in bone scaffolds. In addition, GAGs which are the constituent of bone and cartilage of humans have been shown to have structurally analogous to κ-carrageenan (κ-CG) [17].

The properties of k-carrageenan incorporated HA implants are improved as compared to only HA based implants. Hence, k-carrageenan incorporated HA is likely to achieve the requirements of bioresorbability and biocompatibility for use as scaffolds. However, less number of applications of k-carrageenan have been available that includes micro-encapsulation along with immobilization of certain drugs and wound dressing [18].

Infection has posed to be a major threat in orthopedic implants leading to revision surgery or implant failure. Hence it is a challenging job of treating these infections that may lead to serious complications resulting in amputation and mortality. Infections at the implant sites results in biofilm formation. Because of the higher demand for novel biocompatible materials having antibacterial properties, the researchers started to incorporate antibacterial materials in biomaterials [13, 19]. Ag has got significant importance as an antibacterial agent and has extensively been used in various biomedical applications.

To mimick the natural bone structure, a novel promising approach has been taken in the present study for the fabrication of hydrogels to regenerate the bone and cartilage defect sites simultaneously providing antibacterial efficacy. The sol–gel based methodology was employed to fabricate and maintain the nano-HA precipitates while a lyophilization technique was used to print the structure of the hydrogel.

## 2. Materials and methods

### 2.1 Specimen preparation

nHA was prepared by taking a Ca/P ratio of 1.67 by dropwise addition of 0.6 M calcium hydroxide with 0.4 M orthophosphoric acid and adjusting its pH to 12. The solution was stirred vigorously at 1500 rpm for 12h and were drop wise added to k-carrageenan solution under stirring (10 °C, 20 min) condition. 0.7% Ag(NO_3_)_2_ solution was dropwise added to the above solution. Two separate specimens were prepared with CG concentration of 1.5 wt% and 3 wt%. To obtain homogeneity, samples were stirred at 90 rpm (40 °C, 24 h) and ultrasonicated for 2 h. The nHA-Ag-CG gels were decanted into Teflon molds quenched to −80°C and lyophilized for 36 h.

### 2.2 Swelling studies

The water-absorption capability of the prepared specimens was examined as per equation (1). For measuring the water absorption capability, a known quantity of the lyophilized hydrogel (*Wd*) was immersed in a phosphate buffered saline (PBS) solution at pH 7.4 and 37 °C. The wet hydrogel was taken out of PBS and after wiping off the excessive adsorbed surface PBS, it was weighed to find out the weight during swelling (*Ws*) at different durations (20- and 40-min, 1-, 2-, 4-, 8-, 24-, 48-, and 72-hours). The swelling ratio was measured using the equation

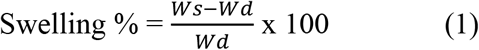

### 2.3 Degradation study

The degradation profile of the prepared specimens was examined as per equation (2). The initial weight (*Wi*) of the lyophilised hydrogels were noted and they were soaked in PBS of pH 7.4. The incubation of hydrogels was carried out at 37 °C and placed in a shaker at 80 rpm. The PBS was renewed every three days. At time points of 3-, 7-, 14-, 21-, and 28-days, the hydrogels were rinsed with water and lyophilized. The dried hydrogels were then weighed (*Wf*) and the degradation profile was measured as per the following equation (2):

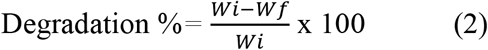

The data presented are the mean ±standard deviation (SD) of five measurements for each hydrogel in swelling and degradation behaviour.

### 2.4 Contact angle measurement

The water wetting angle or hydrophilicity of the hydrogel was quantified by measuring their angle subtended with a drop of PBS at pH 7.4. The contact angle was measured using a Contact angle detection system (Kruss, Germany). The data represented are the mean ± SD of five measurements for each hydrogel [12].

### 2.5 Mechanical properties

Compressive tests were used to study the mechanical properties through two modes such as (i) the hydrogels were compressed till its breaking point and (ii) the hydrogels were placed on cyclic compression test. Initial test was employed to measure the Young’s modulus and compressive strengths while the degradation behavior of hydrogels was estimated from the second test. Hydrogel blocks (8mm diameter x 5 mm thickness, disk shape) were used in both the tests. The solutions of hydrogels were transferred to a 8mm x 5 mm circular pit and irradiated with UV for photocrosslinking. The hydrogel specimens were then dipped in PBS solution for 10 h. The hydrogels in swelling condition were subjected to compression until break using a universal testing machine (UNITEST M2, Test One Co. Ltd., Korea) (load: 100 N, Strain: 5 mm/min, 25 °C) in order to measure their compressive strength and Young’s modulus. Moreover, the durability test was carried out by cyclic compression test as performed earlier. 60% applied load was given for compression and data collected were from 50 cycles. The data were the mean of three tests with SD.

### 2.6 Bacterial viability test

*S. aureus* was taken for the antibacterial efficacy of the specimens. The strain of bacteria was cultured in MH Broth medium [20]. MTT assay was taken to quantify the *in vitro* bacteria viability. The specimens were placed in a 24-multiwell plate each carrying 1 mL bacteria suspension with an amount of 1× 10^6^ CFU/mL and incubated at 37 °C in a CO_2_ atmosphere. The *in vitro* culture of bacteria was performed for 10, 20 and 30 h and the bacteria adhered to the substrate were quantified for bacterial viability by inoculation of equal quantities (200 μL) of bacterial suspension and MTT solution (5 mg/mL) and they were then placed for incubation at 37 °C for 10 h until the formation of formazan crystals. The OD of the specimen was calculated using a spectrophotometric microplate reader at 490 nm [19].

### 2.7 *In Vitro* Cell Culture

Human osteoblast like cells (MG63) were cultured in DMEM added with 10 % fetal calf serum (FCS, HiMedia) and incubated in 5 % CO_2_ at 37 °C. The culture media was renewed every alternate day. After achieving confluence, the non-viable cells were removed by a trypsin–EDTA solution (0.5 g/L trypsin and 0.2 g/L EDTA, Gibco) and the cells were shifted to a new tissue culture flask [21].

### 2.8 Cell Counting Kit-8 (CCK-8) Assay

CCK-8 assay (Dojindo laboratories, Kumamoto, Japan) was used to assess the MG63 cell proliferation of hydrogels. The specimens were placed in a 96-multiwell plate. 1 × 10^4^ cells/well were added on each well and incubated for 1, 7 and 14 days. The specimens were washed thrice with PBS solution to remove the non-viable cells. The specimens were added with 10 µL of CCK-8 solution and incubated for 3 h. The optical density (OD) was then quantified with the help of a microplate reader at 490 nm. For each incubation time of 1, 7 and 14 days, the absorbance values were taken as the mean of three measurements [21].

### 2.9 Osteogenic expressions

The osteoblastic expression of genes such as RUNX2, collagen type I (COL1) and osteocalcin (OC) were assessed using RT-PCR. RNA, extracted with Rneasy® Mini Kit (QIAGEN) as per Instruction manual was measured using UV-VIS spectrophotometry at 260 nm. 0.5 µg of RNA was reverse transcribed and amplified (25 cycles) with RT-PCR at 55 °C. Table 1 depicts the primers used in RT-PCR analyses. ImageJ 1.41 software was used to analyse the bands obtained from the electrophoresis of a 1% (w/V) agarose gel [21].

### 2.10 Confocal laser scanning microscopy (CLSM) analysis

After incubating the specimens for a week, the osteoblast cells were fixed in (4% paraformaldehyde) and then permeabilized by 0.5% Triton X-100. The DAPI (4’,6-diamidino-2-phenylindole, Sigma-Aldrich) fluroscence staining was employed for visualizing nuclei. For observation of osteoblast morphology, images were subjected to CLSM (Leica TCS SP8, Wetzlar, Germany) analysis [19].

## 3. Statistical analysis

The experimental values were reproduced as mean ± standard deviation (SD) and the difference in values of data were reciprocated by one way analysis of variance (ANOVA).

## 4. Results and discussion

Tissue engineering has played a vital role in overcoming major trauma and damaged tissues or organs. Hydrogels have been found to be emerging scaffolds with regenerating impacts on cartilage, soft and hard tissues. Injectable hydrogels have self-healing characteristics that permits the gel to obtain desirable mechanical strength by tissue in-growth [22]. Gelling of hydrogels at pH 7.4 and 37 °C are the emerging biomaterials where gelation occurs due to body temperature without the help of any chemical or mechanical stimuli. Furthermore, these hydrogels can be directly applied in a minimally invasive method [23].

It has long been established that HA and carageenan have regenerative properties as far as vascularization in human bones are concerned. However, because of their higher biocompatible properties they are prone to bacterial infection [24]. Hence, simultaneous presence of an antibacterial agent such as Ag in a hydrogel matrix would provide the dual action of hydrogel in terms of biocompatibility and antibacterial efficacy.

### 4.1 Swelling ratio determination

Swelling is a vital property of hydrogels that stimulates the gas exchange, helps in absorbing body fluid, transferring nutrients, encapsulating and growing cells within the hydrogel. Fig. 1 depicts the swelling ability of hydrogel.

**Fig 1.**
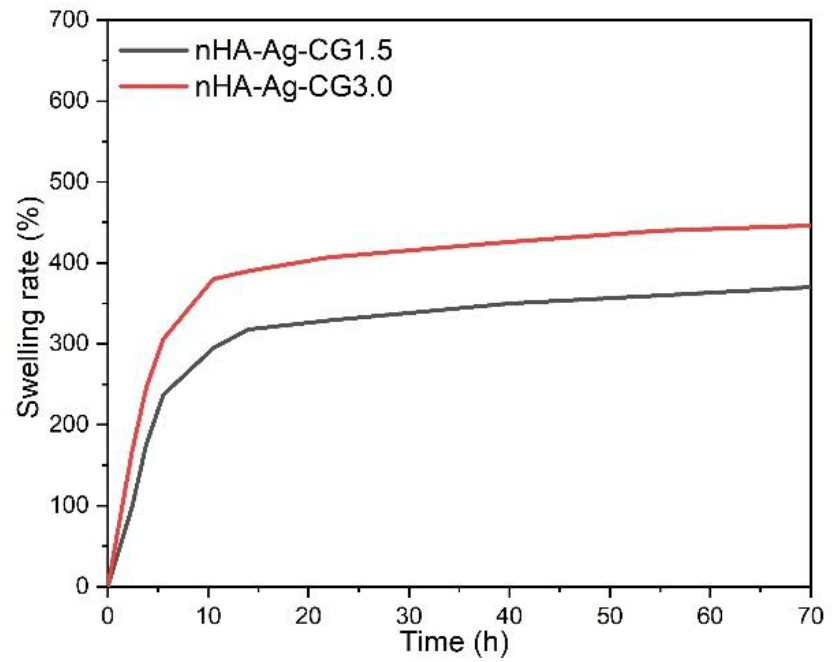
Swelling ability

While nHA-Ag-CG3.0 exhibited higher swelling ratio (500%), it was significantly reduced (400%) in nHA-Ag-CG1.5 hydrogels. The higher swelling ratio is envisaged to induce better cell adhesion and proliferation. Greater swelling index helps in transport of nutrients and fluid uptake from the media near its vicinity. Enhanced swelling ratio may also be due to higher hydrophilicity of carageenan [25]. The swelling equilibrium was attained at 7h for nHA-Ag-CG3.0 while nHA-Ag-CG1.5 attained it at 3 h. From these results, it can be inferred that the incorporation of higher CG concentration in nHA-Ag-CG1.5 hydrogel decreased the swelling ratio. Absorption of body fluid through swelling of the hydrogels is the basic necessity in tissue engineering constructs. In the functions of hydrogels, swelling and water retention ability are primarily related to the amorphous regions and free OH^-^ radicals in the polymer. The swelling rate of the hydrogels should be able to efficiently absorb the biological fluids that come in contact with them. Hence, the evaluation of swelling performance of the hydrogels is a vital criterion.

### 4.2 Biodegradability evaluation

Biodegradability is another vital characteristic of the hydrogels such that once the functional site of the hydrogels is completely regenerated; the scaffold on the other hand has to degrade concurrently. The biodegradability test was performed in PBS at body temperature for 3 weeks. The biodegradation rate of nHA-Ag-CG1.5 was found to be equivalent to nHA-Ag-CG3.0 hydrogels at day 7 while the biodegradation rate increased faster in nHA-Ag-CG3.0 on days 14 and 21 that may be attributed to the possible interaction of nHA and Ag with their CG matrix. The results from swelling and degradation behavior were found to be interrelated. Higher swellable matrices interact with more number of H_2_O molecules that result in quicker degradation of the hydrogels and slowly degrading matrix showed lower swelling ratio [26]. Hence both the results depicted that higher swelling and faster degradation of nHA-Ag-CG3.0 was a consequence of higher CG concentration.

### 4.3 Contact angle measurements

Water wetting angle of the surface of a biomaterial plays a vital role that determines the adhesion of proteins or osteoblast cells on its surface [12]. The hydrophilicity of the hydrogel was obtained by measuring the angle between the hydrogel surface with that of simulated body fluid. It was found from the study that the as-formed hydrogel was hydrophilic in nature with a contact angle of 32°. The water wetting ability of a biomaterial surface largely impacts the cell adhesion, proliferation and differentiation based on their hydrophilicity [27].

### 4.4 Evaluation of mechanical properties

Incorporation of metallic antibacterial agents into polymeric blends at physiological temperature and pH has stiffened the hydrogels. Hydrogels having higher (3%) concentrations of CG showed to have higher tensile strength than the lower (1.5%) concentration of CG. Higher CG concentration also provided higher stiffness (20.67 MPa) than its lower CG (18.79 MPa) counterpart [26, 28]. In the composite hydrogels, increased levels of compressive strength is obtained as compared to hydrogels without inorganic components which is attributed to the reinforcement that develops secondary cross linking points arresting free movement of polymer chain thus making the hydrogels stiff.

The hydrogel scaffolds having adequate load bearing capacity have shown to be ideal as biomedical material. Hence, the test to evaluate the compressive strength for both the hydrogels were performed. Howsoever, the compressive strength was in the order of nHA-Ag-CG3.0 (11.2 MPa) > nHA-Ag-CG1.5 (9.1MPa). It can be found from Fig. 2 and Fig. 3 that the compressive strength and Young’s modulus were considerably higher in nHA-Ag-CG3.0 than nHA-Ag-CG1.5. The Young’s modulus of nHA-Ag-CG1.5 is 0.36 ± 0.07 MPa that increased in case of nHA-Ag-CG3.0 (0.54 ± 0.09 MPa) demonstrating better interfacial compatibility of their matrix with respect to the reinforcement. This increase in reinforcement concentration results in higher stiffness that dissipate energy. Hence, it can be found from the results that nHA-Ag-CG3.0 improved its mechanical properties after incorporation of higher CG concentration. As the hydrogels are stressed, the entangled chains of polymeric matrix receive the load and gets reoriented resulting in draining out of interstitial fluid. Scaffold structure with adequate mechanical strength plays an important role for a successful implant application. Higher mechanical strength in both the hydrogels may be attributed to the homogeneous distribution of hydroxyapatite particles.

**Fig 2.**
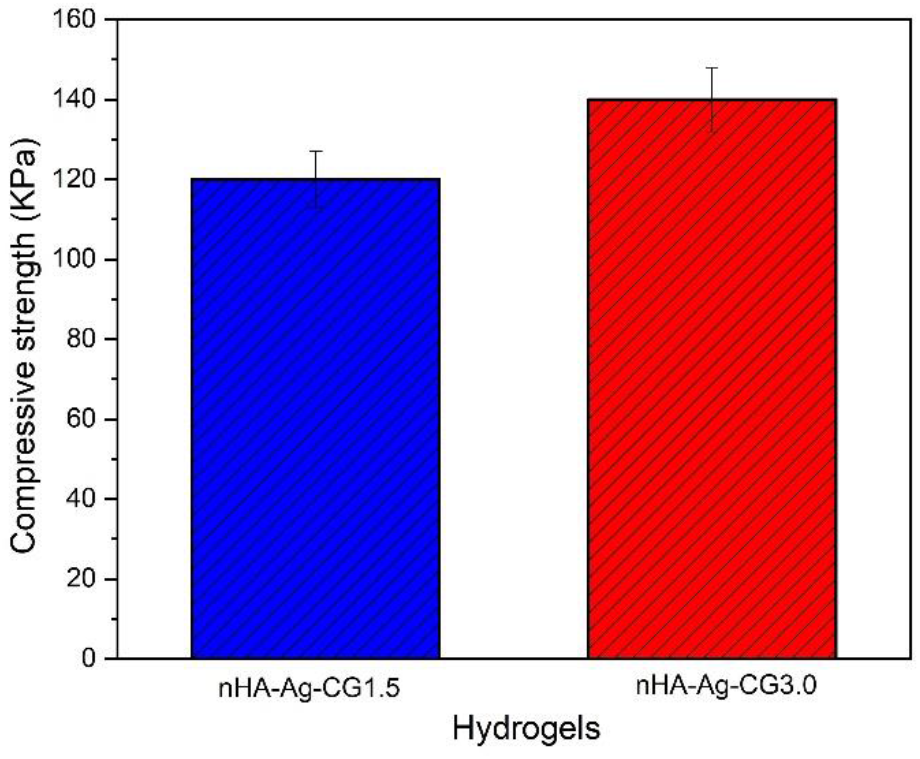
Compressive strengths of the specimens

**Fig 3.**
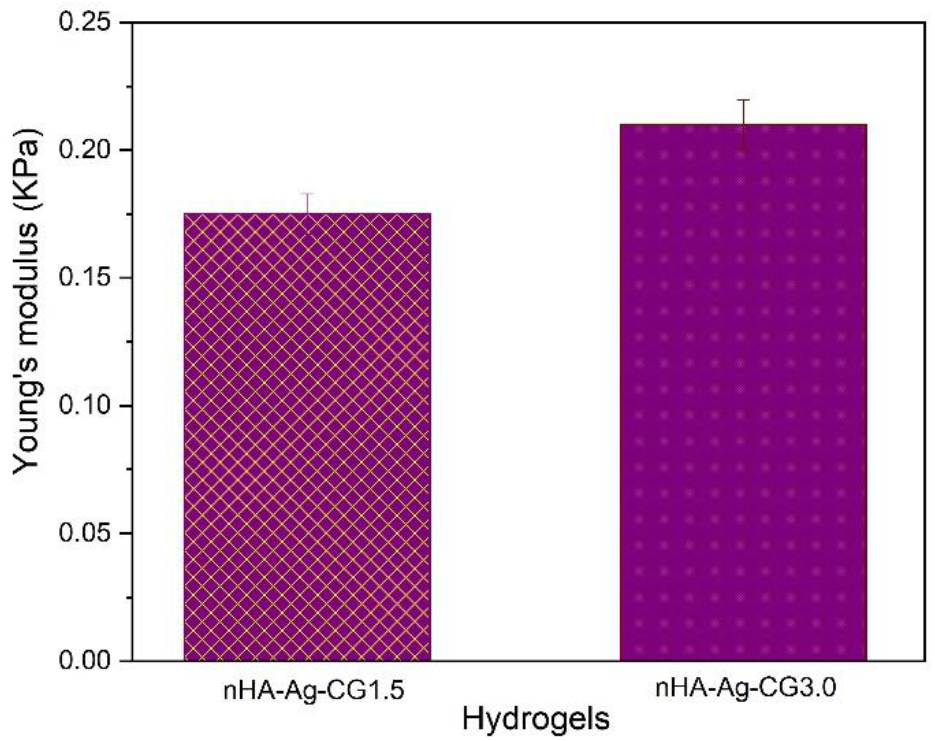
Young’s modulus of the specimens

### 4.5 Antibacterial activity

In order to assess the antibacterial efficacy of Ag present in the hydrogels, their antibacterial activities were evaluated using *S. aureus* pathogenic bacteria. While the concentration of Ag was same in both the hydrogels, the antibacterial activities of both of the hydrogels were expected to be equivalent. Hence, a comparison of the antibacterial activities of nHA-Ag-CG3.0 hydrogel with that of nHA-Ag-CG1.5 hydrogels is depicted in Fig. 4. Antibacterial Ag blocked the channels of bacteria cells and the positive charge of Ag neutralize the negatively charged bacteria membrane changing the lipid layer structure, increasing the permeability of cell membranes and ruptured the bacterial membrane resulting the bacteria cell death [13, 29]. The bacterial cell viability of *S. aureus* was performed after 10 h, 20 h, and 30 h of culture. The nHA-Ag-CG3.0 exhibited restrained growth of *S. aureus* as compared to nHA-Ag-CG1.5 at 30 h because of the presence of higher concentration of biocompatible CG.

**Fig 4.**
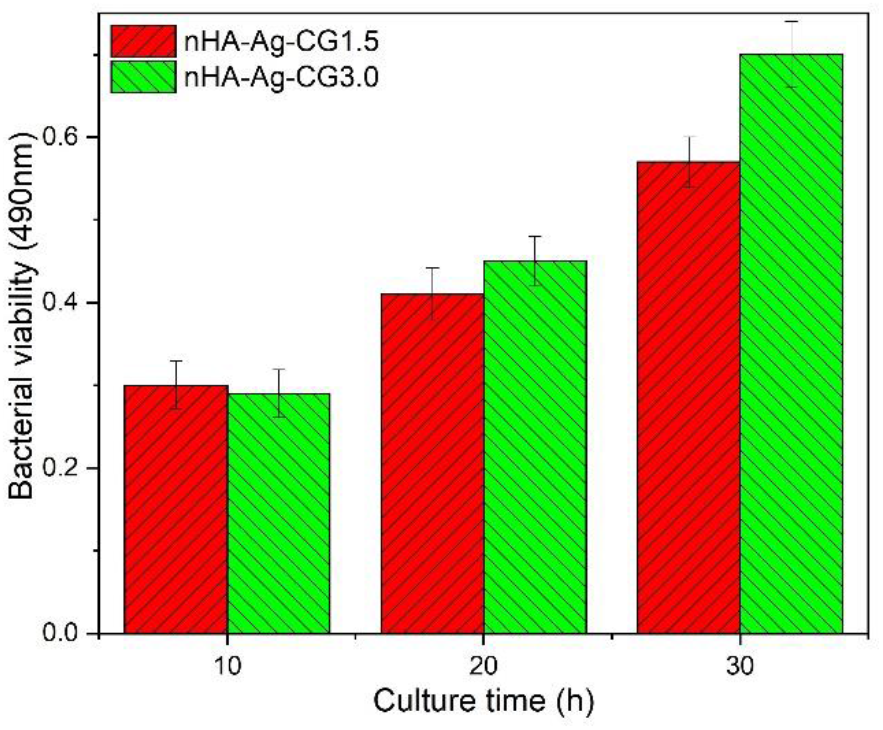
Bacterial viability on the specimens

### 4.6 Cell viability assay

The osteoblast cell viability of both the as-formed hydrogels was performed using CCK-8 assay as depicted in Fig. 5. It can be seen that higher MG63 cells were survived on the nHA-Ag-CG3.0. This may be attributed to the presence of higher amount of CG in the hydrogels. It has been established that CG is responsible for higher Ca deposition because of its affinity to bind Ca that subsequently accelerate cell differentiation giving rise to new bone formation [17].

**Fig 5.**
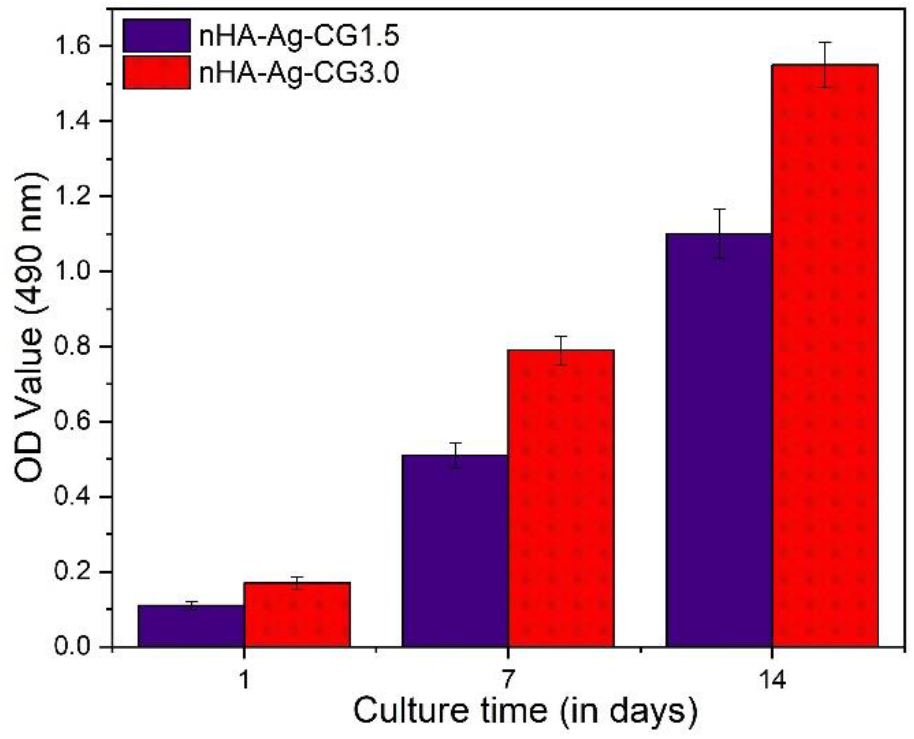
MG63 cell viability on the specimens

Hence, nHA-Ag-CG3.0 may be considered to have more cytocompatibility than nHA-Ag-CG1.5. Although the cytocompatibility of the hydrogels is compromised in the presence of antibacterial agent, presence of cytocompatible HA and CG compensate the cytotoxic behavior of Ag. These hydrogels are suggestive of being used as successful scaffold materials for hosting bone tissues.

The qualitative analysis of MG63 cell attachment on the hydrogels was performed using CLSM imaging after 24 h of culture. DAPI staining was used to visualize nucleus of adherent cells on the hydrogels. It can be found that cell spreading and proliferation were higher in nHA-Ag-CG3.0 hydrogels after 12h of culture. This may be attributed to the higher cytocompatibility of nHA-Ag-CG3.0. Hence, the as-formed hydrogels may be useful in contributing to osteogenicity in the presence of antibacterial agents solving a long standing problem of dual action.

Osteogenic gene expression studies demonstrated that RUNX2 (Fig. 6) and OC (Fig. 7) expressions for nHA-Ag-CG3.0 were higher by 29% and 27% respectively as compared to nHA-Ag-CG1.5 while there was 40% increase in COL1 (Fig. 8) expression on day 1. On the other hand, RUNX2 and OC expressions for nHA-Ag-CG3.0 were higher by 38% and 28% respectively as compared to nHA-Ag-CG1.5 while there was 45% increase in COL1 expression on day 7. Moreover, RUNX2 and OC expressions for nHA-Ag-CG3.0 were higher by 29% and 45% respectively as compared to nHA-Ag-CG1.5 while there was 36% increase in COL1 expression on day14. The steady increase in osteogenic expressions for nHA-Ag-CG3.0 specimen may be attributed to the presence of higher concentration of carageenan that contributed for higher osteogenicity. COL-I osteogenic marker is an early marker exhibiting mineralization of bone tissues while other marker RUNX2 is expressed in the later stages during bone mineralization while OC is expressed during remodeling phase [30]. nHA-Ag-CG3.0 promoted the expression of COL-I, RUNX2 and OC markers exhibiting exceptional bioactivity. However, further experiments may be conducted to explain the complete signaling pathways and the intrinsic mechanisms involved.

**Fig 6.**
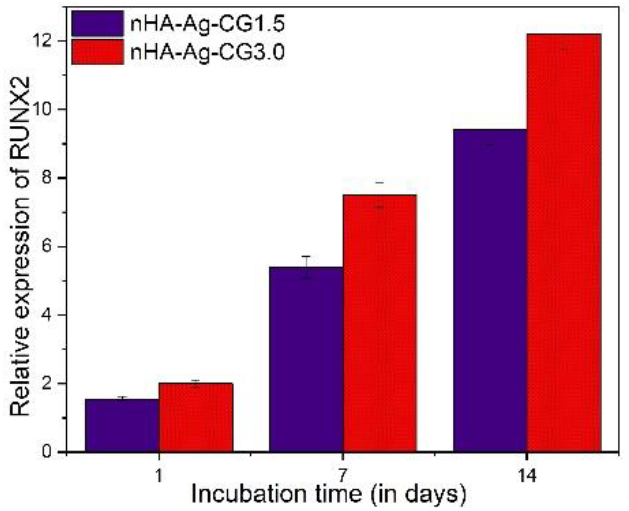
Relative expression of RUNX2

**Fig 7.**
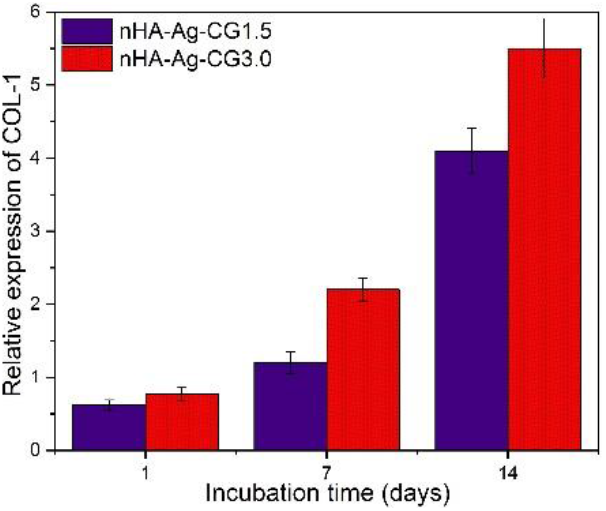
Relative expression of COL-1

**Fig 8.**
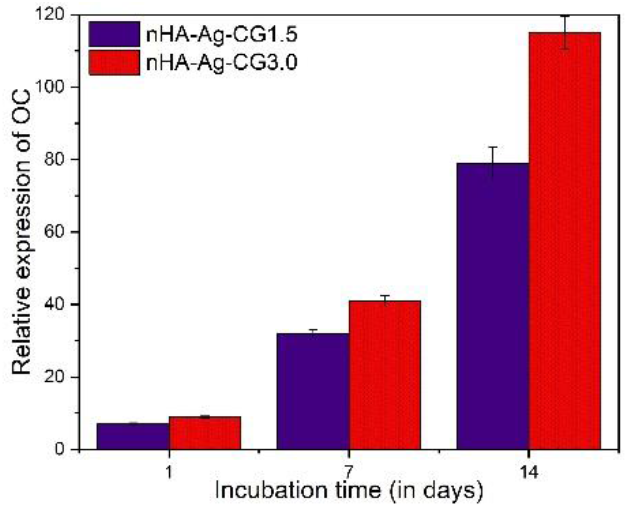
Relative expression of OC

Increased surface area has been shown to impact more on the cells. The polar surface of nHA is largely hydrophilic similar to carageenan that helps in the adsorption of protein based on the principles of ionic and hydrogen bonding. Hydrophilic surfaces induce cellular activities and has been shown to stimulate vitronectin and fibronectin those in turn supports the growth of osteoblast cells protecting the bioactivity of adsorbed protein. It has been found from many studies that Ca^2+^ and PO_4_^3-^ of HA and SO_4_ ^2-^ of CG helps in promoting adsorption of many proteins [31].

### 4.7 CLSM analysis

CLSM analysis (Fig. 9) of the specimens at 6 h of culture depicted that osteoblasts adhered to the surface of specimens and offered circular morphology on the nHA-Ag-CG1.5 specimen and elongated morphology on the nHA-Ag-CG3.0 specimen at 6 h of culture. Higher magnification of CLSM images indicated distinct differences in the morphology of cells in both the specimens and visibility of extended F-actin cytoskeletal structure of cells on nHA-Ag-CG3.0 specimen. On the nHA-Ag-CG1.5 specimen, MG63 cells offered low cytoplasm expansion in distinctive osteoblast cells with rounded morphology with clear visibility of the nuclei. Surveillance of MG63 cells at 6 h of culture offered evidence of cell adhesion on the surfaces of specimens and on the cytoplasm expansion and the attainment of distinctive cell morphology, actions which are strongly reliant on the reform of the F-actin cytoskeleton. It is an important observation since the F-actin cytoskeleton takes primary role in the maintenance of the shapes and junctions of cells that provides mechanical backing to the cells.

**Fig 9.**
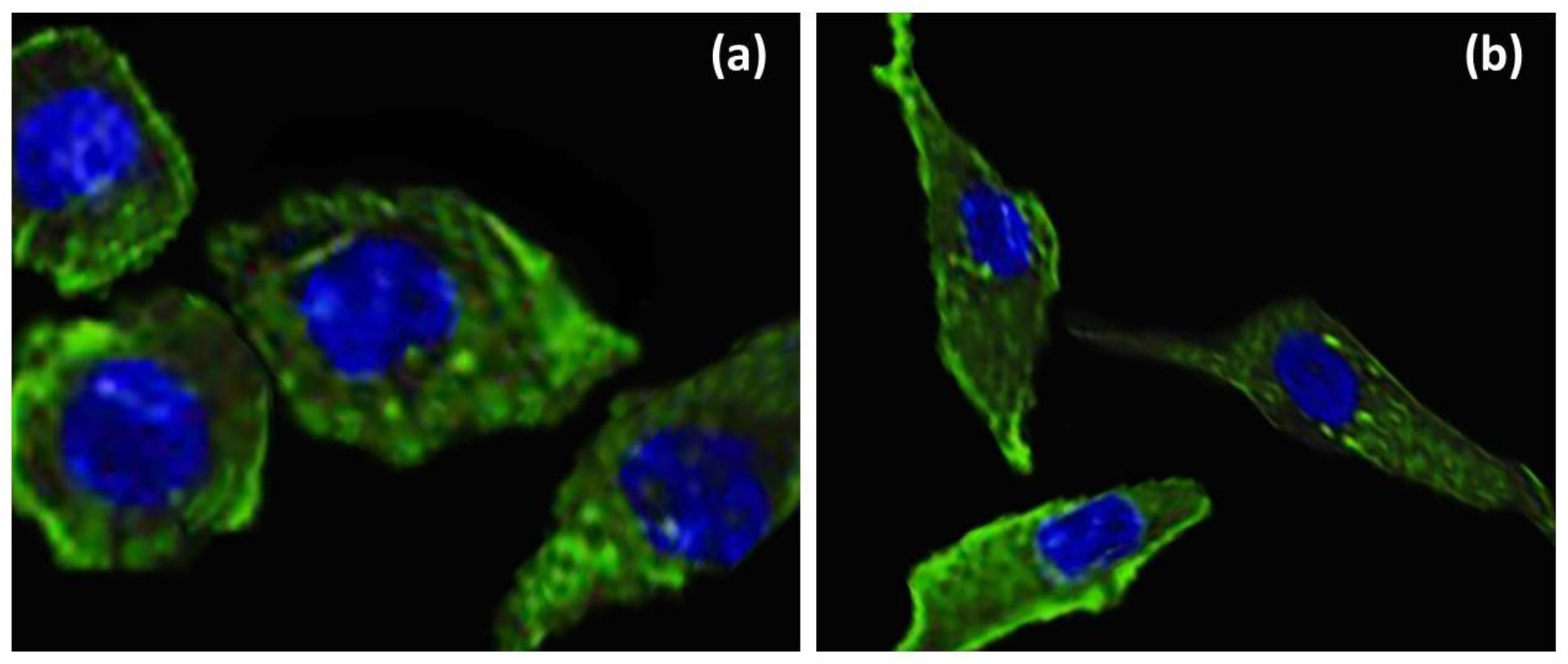
CLSM images of (a) nHA-Ag-CG1.5 and (b) nHA-Ag-CG3.0

## 5. Conclusions

The as-formed hydrogels were hydrophilic in nature. nHA-Ag-CG3.0 hydrogel improved its mechanical strength after incorporation of higher CG concentration. The Young’s modulus of nHA-Ag-CG3.0 was higher than nHA-Ag-CG1.5 demonstrating better interfacial compatibility of their matrix with respect to the reinforcement. The higher swelling ratio and biodegradability of nHA-Ag-CG3.0 is envisaged to induce better cell adhesion and proliferation. The bacterial cell viability of *S. aureus* on nHA-Ag-CG3.0 showed to have restrained growth as compared to nHA-Ag-CG1.5. nHA-Ag-CG3.0 may be considered to have more cytocompatibility than nHA-Ag-CG1.5 due to higher concentration of CG. Osteogenic gene expression studies demonstrated that nHA-Ag-CG3.0 significantly elevated the expression of osteogenic genes such as COL1, RUNX2 and OC during the culture times of 1, 7 and 14 days. Incorporation of higher CG amount in the presence of Ag was found to be more cytocompatible with better mechanical properties to be used as an ideal scaffold material.

